# PIMBA: a PIpeline for MetaBarcoding Analysis

**DOI:** 10.1101/2021.03.23.436646

**Authors:** Renato R. M. Oliveira, Raíssa Silva, Gisele L. Nunes, Guilherme Oliveira

**Affiliations:** Instituto Tecnológico Vale, Belém, Pará, Brazil; Programa de Pós-Graduação em Bioinformática, Instituto de Ciências Biológicas, Universidade Federal de Minas Gerais, Belo Horizonte, Minas Gerais, Brazil

## Abstract

**Summary:** DNA metabarcoding is an emerging monitoring method capable of assessing biodiversity from environmental samples (eDNA). Advances in computational tools have been required due to the increase of Next-Generation Sequencing data. Tools for DNA metabarcoding analysis, such as MOTHUR, QIIME, Obitools, and mBRAVE have been widely used in ecological studies. However, some difficulties are encountered when there is a need to use custom databases. Here we present PIMBA, a PIpeline for MetaBarcoding Analysis, which allows the use of customized databases, as well as other reference databases used by the softwares mentioned here. PIMBA is an open-source and user-friendly pipeline that consolidates all analyses in just three command lines.

**Availability and Implementation:** https://github.com/reinator/pimba

## Introduction

DNA metabarcoding is a powerful tool that has been widely used for biodiversity monitoring and ecosystem assessment from DNA information. The technique based on the high-throughput sequencing (HTS) allows the multispecies detection with DNA from environmental samples (eDNA) using a specific molecular marker for specific groups (plants, animals, fungi, and bacteria) (Creer *et al*., 2016). Next-Generation Sequencing (NGS) results in millions of DNA sequences (reads) that are sequenced to decipher the genetic code of species that help to answer taxonomic and functional questions. For DNA metabarcoding analysis, the reads are grouped into operational taxonomic units (OTUs) to differentiate species or taxa based on the similarity of genetic code(Cristescu, 2014; Hering *et al*., 2018). Similarity of 97% is commonly used as a cutoff, but this value depends on the group to be evaluated (Deiner *et al*., 2017).

The recent increase of NGS data has caused the development of new tools for DNA metabarcoding analysis, making the metabarcoding method more accessible and user-friendly (Alberdi *et al*., 2018). Mothur (Schloss *et al*., 2009), Qiime (Caporaso *et al*., 2010), Obitools (Boyer *et al*., 2016), and mBRAVE (Ratnasingham, 2019) are currently the most used tools for metabarcoding analysis. The biggest restriction of these pipelines is to make it difficult to use a customized database for taxonomy assignment. For example, Mothur is useful when analyzing 16S/18S rRNA, Influenza viral, and fungal ITS regions, using Greengenes (DeSantis *et al*., 2006), Influenza Virus, and SILVA (Quast *et al*., 2013) databases, respectively. Qiime (and even its updated version, Qiime2) is optimized to analyze metabarcoding data from 16S rRNA, 18S, and fungal ITS marker genes, using Greengenes, SILVA, and UNITE (Abarenkov *et al*., 2010) databases, respectively. Qiime2 allows the user to train a classifier with a Na-iveBayes model, but they report tests by using only a 16S example and with some constraints to the use of this classifier with other marker genes. Qiime does not support adapting its pipeline to use a custom database to analyze sequenced data from different marker genes, such as COI or Plant ITS. Obitools is optimized to analyze data from 16S (SILVA and PR2) and it also allows the use of the NCBI database for taxonomic assignment. mBRAVE is optimized to use only the BOLD database as a reference, allowing the researcher to use a personalized database only after BOLD submission.

We developed PIMBA, a PIpeline for MetaBarcoding Analysis, which adapts the Qiime pipeline for OTUs clustering with additional OTU corrections based on the algorithm LULU (Frøslev *et al*., 2017). A preliminary abundance and diversity analysis are also automatically delivered. The main innovation of this pipeline is the ease of using both standard and customized databases minimizing errors in taxonomic assignments making the results more reliable.

## Implementation

Besides implementing all the features provided by the other metabarcoding tools, PIMBA also allows the user to apply different databases and not only those commonly used by most of the available softwares. PIMBA can be used with single or paired-end reads and is divided into three steps: (1) preprocessing, which promotes the demultiplexing and quality treatment of reads (Figure 1A); (2) clustering, in which reads are clustered into OTUs, along with errors correction and taxonomic assignment (Figure 1B), and (3) plotting, in which alfa and beta diversity plots are built by Phyloseq (McMurdie and Holmes, 2013), including rarefaction curves and Principal Coordinates Analysis (PCoA), respectively. A metadata file is required for this last step.

**Fig. 1:**
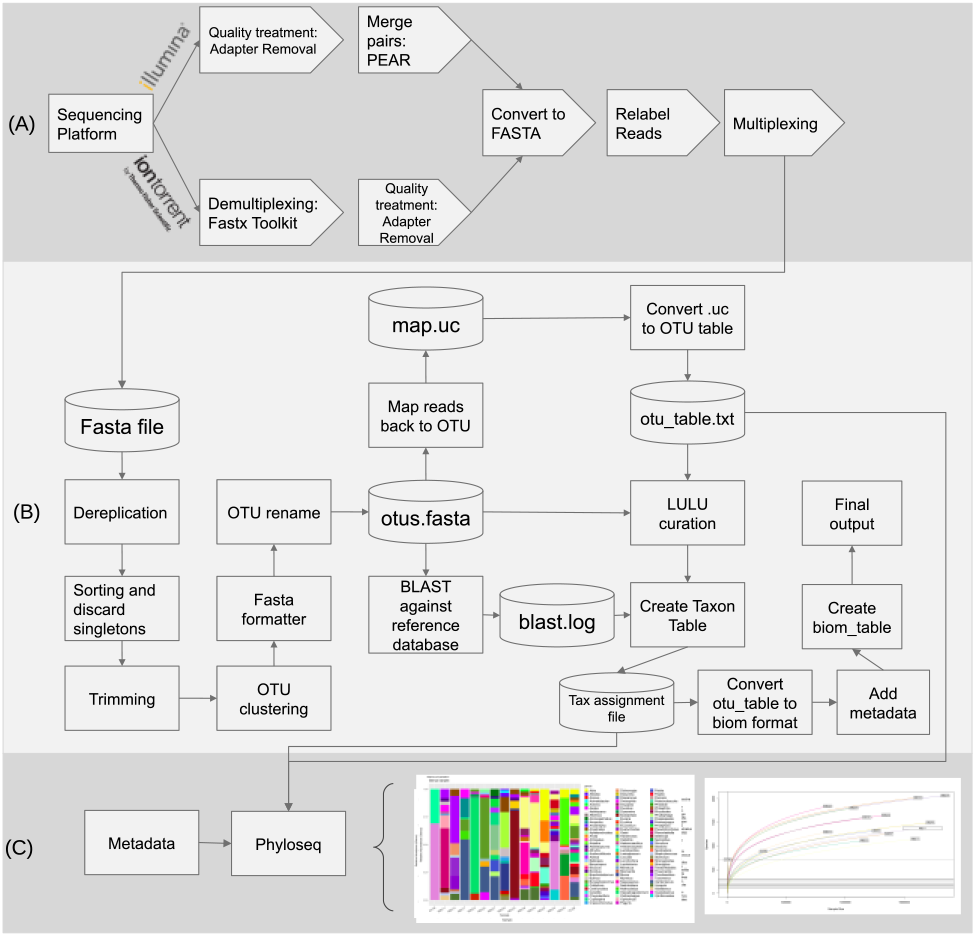
Workflow of PIMBA steps for preprocessing (1A), clustering (1B) and plotting (1C).

With only three command lines, PIMBA runs the metabarcoding analysis, allowing the user to easily change parameters, including OTU similarity and length, assignment similarity and coverage, number of threads to use, etc.). PIMBA also allows the user to analyze datasets from distinct molecular markers (COI, Plant ITS, rbcL, matK, etc.) from unpublished databases. Thus, it is only needed a fasta file with the reference sequences and their identification, and a two-column tax.txt file with the sequence ID and the full taxonomy written for every reference sequence in the fasta file. In the case of reference databases such as UNITE, SILVA, and GREENGENES, the format used is the same used by QIIME, and the user will only need to configure the paths to those reference databases in the databases.txt file. The pipeline and user guidance are available for download on GitHub (https://github.com/reinator/pimba).

## Results

To demonstrate that PIMBA is effective at analyzing a metabarcoding dataset, we performed a benchmark using a simulated amplicon sequencing from the 16S marker gene “extracted” from a soil environment (Almeida *et al*., 2018). The dataset comprises 100 species from 70 genera commonly found in soil samples with a similar relative abundance.

The table below shows the accuracy of PIMBA when using SILVA 16S (which uses the same steps as QIIME) or the NCBI database.

**Table 1.** Benchmark results for using PIMBA with SILVA 16S or NCBI database.

| Database | Number of OTUs | PIMBA's sensibility |
| --- | --- | --- |
| SILVA | 100 | 85% |
| NCBI | 100 | 85% |

The sensibility results indicate how much of the taxa containing in the simulated dataset was obtained by PIMBA using the different databases.

Results show that whether using SILVA or the NCBI database, PIMBA can obtain the same sensibility when identifying taxa in a simulated database. Since for the SILVA database the steps were the same that QIIME uses, results also show that the efficiency of both PIMBA and QIIME are the same for this simulated dataset.

We also used PIMBA to analyze two metabarcoding datasets (Plant ITS and Invertebrates COI). PIMBA generated all the outputs needed to infer results and additional metabarcoding analysis, as well as alfa and beta diversity analyses, PCoA, and taxonomic bar plots.

